# Targeting Scavenger Receptor Type B1 In Cholesterol-Addicted Lymphomas Abolishes Glutathione Peroxidase 4 and Results in Ferroptosis

**DOI:** 10.1101/2020.06.16.155242

**Authors:** Jonathan S. Rink, Adam Yuh Lin, Kaylin M. McMahon, Andrea E. Calvert, Shuo Yang, Tim Taxter, Jonathan Moreira, Amy Chadburn, Amir Behdad, Reem Karmali, C. Shad Thaxton, Leo I. Gordon

## Abstract

Normal human cells can either synthesize or uptake cholesterol from lipoproteins to meet their metabolic requirements. Some malignant cells absolutely require cholesterol uptake from lipoproteins for survival because *de novo* cholesterol synthesis genes are transcriptionally silent or mutated. Recent data suggest that lymphoma cells dependent upon lipoprotein-mediated cholesterol uptake are also dependent on the expression of the lipid hydroperoxidase enzyme glutathione peroxidase 4 (GPX4) to prevent cell death by ferroptosis. Ferroptosis is an oxygen-and iron-dependent cell death mechanism that results from the accumulation of oxidized lipids in cell membranes. To study mechanisms linking cholesterol uptake with ferroptosis, we employed lymphoma cell lines known to be sensitive to cholesterol uptake depletion and treated them with high-density lipoprotein-like (HDL) nanoparticles (HDL NPs). HDL NPs are a cholesterol-poor ligand of the receptor for cholesterol-rich HDL, scavenger receptor type B-1 (SCARB1). Our data reveal that HDL NP treatment activates a compensatory metabolic response in treated cells favoring *de novo* cholesterol synthesis, which is accompanied by reduced expression of GPX4. As a result, accumulation of oxidized membrane lipids leads to cell death through a mechanism consistent with ferroptosis. Furthermore, ferroptosis was validated *in vivo* after systemic administration of HDL NPs in mouse lymphoma xenografts and in primary samples obtained from patients with lymphoma. In summary, targeting SCARB1 with HDL NPs in cholesterol uptake addicted lymphoma cells abolishes GPX4 and cancer cell death ensues through a mechanism consistent with ferroptosis.

## INTRODUCTION

Despite long-term remission observed in some patients with lymphoma, greater than one third of patients with the most common subtype, diffuse large B cell lymphoma (DLBCL), will relapse or have disease that is refractory to primary treatment (1–3). This is especially the case for patients in high-risk groups identified by molecular and clinical prognostic factors (4,5). Experimental therapies for these patients, including immunotherapy and cell-based therapies, have modest success rates, high cost, and toxicity (6,7). Thus, there is a significant need for a new therapeutic approach.

Recently, DLBCL was identified as a cancer type particularly sensitive to cell death by ferroptosis (8). Ferroptosis is an oxygen-and iron-dependent form of necroptosis characterized by accumulation of cell membrane lipid and cholesterol peroxides that results from the targeted inhibition of the lipid hydroperoxidase glutathione peroxidase 4 (GPX4) (9–11). Cells become vulnerable to ferroptosis after GPX4 inhibition because the enzyme reduces and detoxifies lipid peroxides (L-OOH) by converting them to corresponding lipid alcohols (L-OH) (8,10). Malignant cells under oxidative stress are significantly more sensitive to ferroptosis because of higher levels of reactive oxygen species and a reliance on GPX4 activity to mitigate toxic L-OOH accumulation (12–14). This has driven significant interest in the development of treatments to induce ferroptosis in cancer cells. Small molecule inhibitors of GPX4 have been developed and tested, but they are toxic and lack specificity (15), which limits *in vivo* use and clinical relevance.

Data suggest a link between cellular cholesterol uptake and ferroptosis. For instance, removal of lipoproteins from media used to culture ALK^+^ anaplastic large T cell lymphoma (ALK^+^ ALCL) cell lines (SR-786, SUDHL1) and a histiocytic lymphoma cell line (U937) induced ferroptosis (16). The data focused on the role of cholesterol-rich low-density lipoprotein (LDL) binding to the LDL receptor (*LDLR*) as responsible for cholesterol delivery to the malignant cells. Analyses of these cell lines revealed that a loss of squalene epoxidase expression (*SQLE*, aka squalene monooxygenase, ALK^+^ ALCL cells), through hypermethylation of its promoter region, or 3-ketosteroid reductase expression (*HSD17B7*, U937 cells), through a mutational defect, rendered the cells auxotrophic for cholesterol. SQLE and HSD17B7 are rate-limiting enzymes in *de novo* cellular cholesterol biosynthesis, which explains the cellular obligation for cholesterol uptake. Inhibition of LDL-mediated cholesterol uptake by reduction of the LDLR reduced the viability of ALK^+^ ALCL cells and ALK^+^ ALCL patient derived xenografts (PDX) *in vivo,* and sensitized the cells to ferroptosis by GPX4 inhibitors. These data suggest that reduction of cellular cholesterol uptake renders cells sensitive to ferroptosis; however, the mechanism remains unclear.

Scavenger receptor type B-1 (SCARB1) is a high-affinity receptor for cholesterol-rich high-density lipoproteins (HDL), which has been implicated as a target in human cancers based upon its role in cancer cell cholesterol uptake (17–25) and in membrane-anchored second messenger signaling pathways (21,26–29), among other factors (30–33). Our group developed cholesterol-poor high-density lipoprotein (HDL)-like nanoparticles (HDL NPs) to target and tightly bind SCARB1 to prevent the cellular uptake of cholesterol ester from HDLs (21,24,34). We have demonstrated that certain B cell lymphomas highly express SCARB1, and targeted treatment with the HDL NPs potently induces *in vitro* and *in vivo* cell death by inhibiting cholesterol ester uptake and reducing cell cholesterol (21,24,25). We have shown that HDL NP-mediated inhibition of cholesterol ester uptake in highly sensitive B cell lymphoma cells, including Burkitt’s lymphoma (e.g. Ramos cell line) and DLBCL (e.g. SUDHL4 cell line), is accompanied by compensatory increases in the expression of genes required for *de novo* cholesterol synthesis (21). In fact, our data showed lymphoma cells not initially sensitive to HDL NP-mediated cholesterol depletion, including activated B cell (ABC) lymphoma (e.g. TMD8 and HBL-1 cell lines), exhibit a high baseline expression of *de novo* cholesterol biosynthesis genes and higher cholesterol content relative to the highly sensitive lymphoma cell lines (21). These data are consistent with the report by Chen et al showing that more active second messenger pathways downstream of the B cell receptor (BCR) results in *de novo* cholesterol biosynthesis (35). In the case of the ABC lymphoma cells, our data show that cell death can be induced with a synergistic combination of HDL NP and targeted inhibitors of downstream BCR tyrosine kinases (21). Thus, our data show that targeting of SCARB1 by HDL NP with reduction of cell cholesterol uptake and compensatory increase in *de novo* cholesterol biosynthesis is a potential translational model for treatment of cholesterol uptake dependent lymphoma.

With this background our group explored whether or not HDL NP therapy targeting SCARB1 induced lymphoma cell death through a mechanism involving GPX4 and ferroptosis. Initially, we employed a gene microarray as an unbiased approach to study changes in gene expression caused by HDL NPs in a cholesterol uptake dependent lymphoma cell line. These data revealed that HDL NPs obligate cellular expression of *de novo* cholesterol biosynthesis genes, which is accompanied by reduced GPX4 expression. We show herein that reduced GPX4 expression leads to an increase in membrane oxidized lipids and cell death through a mechanism consistent with ferroptosis in cell lines, in an *in vivo* xenograft model, and in primary samples obtained from patients with lymphoma.

## MATERIALS AND METHODS

### Cell Lines

The Ramos (RRID: CVCL_0597), SUDHL4 (CVCL_0539), Raji (CVCL_0511), Daudi (CVCL_0008), SUDHL6 (CVCL_2206), Namalwa (CVCL_0067), Jurkat (CVCL_0367), SUDHL1 (CVCL_0538), SR-786 (CVCL_1711), and U937 (CVCL_0007) human cell lines were obtained from ATCC and were used within 3 months of receipt and/or resuscitation. ATCC uses short tandem repeat (STR) profiling to authenticate their cell lines prior to shipping. For SUDHL4 cells, Charles River Laboratories was contracted to test for mycoplasma contamination prior to use in animal experiments. All cell lines were cultured in RPMI 1640 supplemented with 10% fetal bovine serum (FBS) and 1% penicillin/ streptomycin at 37°C in a humidified, 5% CO_2_ incubator.

### HDL NP Synthesis

The HDL NPs were synthesized and quantified as previously described (36). 5nm diameter citrate stabilized gold nanoparticles (AuNP) were surface-functionalized with apolipoprotein A-I, followed by addition of the phospholipids, 1,2-dipalmitoyl-*sn*-glycero-3-phosphoethanolamine-N-[3-(2-pyridyldithio)propionate] (PDP PE) and 1,2-dipalmitoyl-*sn-*glycero-3-phosphocholine (DPPC). The HDL NPs were purified using the KrosFlo TFF (Tangential Flow Filtration) system with a 50kDa cut-off PES module. The concentration of HDL NPs was calculated using UV-Vis spectroscopy and Beer’s law.

To synthesize fluorescently labeled HDL NPs, the intercalating dye DiI (1,1’-Dioctadecyl-3,3,3’,3’-tetramethylindocarbocyanine perchlorate) was added at a 1µM final concentration during the phospholipid addition step. Purification and quantification of the fluorescently labeled HDL NPs was conducted, as described above.

### HDL NP Binding to SCARB1 Assay

Ramos, SUDHL4, and Jurkat cells were incubated with DiI HDL NPs (10nM) in standard culture media for 2 hours at 37°C, in the presence or absence of the SCARB1 blocking antibody (Novus Biologicals; 1:100; RRID: AB_1291690), and/or the rabbit IgG isotype control antibody (Novus Biologicals; 1:100). Cells were washed once with 1mL of ice-cold FACS buffer (PBS, 1% bovine serum albumin, 0.1% sodium azide) and re-suspended in 500µl of ice-cold FACS buffer prior to flow cytometric analysis (BD LSR II Fortessa). Data were analyzed using the FCS Express software.

### Microarray Analysis

Ramos cells were treated with HDL NPs (40nM), human HDL (40nM) or PBS for 48 hours prior to RNA isolation using the RNeasy Mini kit (Qiagen). RNA samples were converted to cDNA libraries by the Northwestern University Genomics Core facility and were then run on the Illumina HT-12 microarray. A total of 3 biological replicates were run for each condition. Data were analyzed by the Genomics Core facility, with a fold change of >1.5 or <-1.5 and a p value < 0.05 considered significant. Microarray data are available at NCBI GEO under accession number GSE98028.

### Western Blot Analysis

Western blots were conducted as previously described (21). Blots were imaged using the Azure 3000 imager. The SCARB1 antibody (Abcam, RRID: AB_882458; 1:1,000), the GPX4 antibody (Abcam, AB_941790; 1:5,000), the β-actin antibody (Cell Signaling Technologies, AB_2223172; 1:3,000) and a secondary antibody (goat anti-rabbit HRP, Bio-Rad, AB_11125142; 1:2,000) were used in these experiments.

### RT-qPCR Analysis

Ramos, SUDHL4, SUDHL1, SR-786 and U937 cells were treated with HDL NPs (20nM, 50nM), human HDL (hHDL; 50nM) or PBS for up to 72 hours, and RNA isolated using the RNeasy Mini kit (Qiagen). In all cases, hHDL was added at an equimolar concentration to HDL NPs based upon protein concentration, as previously described (24). RNA samples (500ng RNA/ 30µl reaction) were reverse transcribed using a TaqMan Reverse Transcription kit, and qPCR was performed using Taqman Gene Expression Assays (Life Technologies) on a BioRad CFX-Connect iCycler. Samples were standardized to β-actin, and relative expression was calculated using the ΔΔCt method. Biological triplicates were run for each condition.

### C11-BODIPY Assay for Lipid Peroxidation

Ramos, SUDHL4, SUDHL1, SR-786 and U937 cells (2.5 × 10^5^ cells/ ml) were treated with HDL NPs (50nM) or PBS for 24, 48 or 72 hours. Following treatment, C11-BODIPY (1µM final concentration; Thermo Fisher Scientific) was added to each well and the cells were incubated for 30 minutes at 37oC, 5% CO_2_. The cells were then washed twice with 1 X PBS, re-suspended in ice-cold FACS buffer and C11-BODIPY fluorescence in the FITC channel quantified using the BD LSR II Fortessa flow cytometer. Data were analyzed using the FCS Express software.

### Cell Death (MTS) Assay

MTS assays (CellTiter; Promega) were conducted as described previously (21,24). For the Ramos, SUDHL4, Raji, Daudi, Namalwa, SUDHL6 and Jurkat cells were plated at a density of 2 × 10^5^ cells/mL and cultured for 72 hours prior to assay. SUDHL1, SR-786, and U937 cells were plated at a density of 5 × 10^4^ cells/ mL and cultured for 5 days prior to MTS assay. The SCARB1 blocking and isotype control antibodies were added at a dilution of 1:1000 to 1:250. Ferrostatin-1 and deferoxamine (DFO) were obtained from Sigma Aldrich, and added at a final concentration of 1µM. MTS values were standardized to PBS control.

### Tumor Xenograft Model

All animal work was conducted in accordance with Northwestern University’s IACUC and CCM facilities under an approved animal protocol (NU IACUC IS00002415). SCID-beige mice (4 to 6 weeks old; Charles River) were used for the SUDHL4 tumor xenograft study. Flank tumors were initiated using 1 × 10^7^ SUDHL4 cells per mouse. Tumors were allowed to reach ~100mm^3^ before HDL NP treatments began. Based on their initial tumor volumes, mice were randomly divided into 2 groups, PBS (100μL) and HDL NPs (100μL of 1μM NPs). Treatments (intravenous) were administered 3 times per week for 1 week. Tumors were then harvested, and single cell suspensions generated by mechanically dissociating the tumors and passing the cells through a 70-micron filter. C11-BODIPY (1µM final concentration) was added to a fraction of the resultant cell suspension (1 × 10^6^ cells) and flow analysis was carried out as described above. RNA was isolated from the remainder of the cells to quantify GPX4 expression by RT-qPCR, as described above.

### Human Tissue Analysis

Archived, formalin-fixed, paraffin embedded tissue sections were analyzed from patients with large B cell lymphoma and follicular lymphoma. All samples were de-identified of all information other than final diagnosis. A total of 104 follicular lymphoma and 49 diffuse large B cell lymphoma archival samples were obtained and stained for SCARB1 expression. Immunohistochemical staining of the sections was performed using a monoclonal SCARB1 antibody (Abcam, AB_882458; 1:100 dilution) by the Pathology Core at the Robert H. Lurie Comprehensive Cancer Center of Northwestern University. Liver and thymus specimens were utilized as positive and negative controls, respectively. Bright field images were captured at 10X and 40X magnifications.

### Primary Lymphoma Cell Isolation and Analysis

Primary lymphoma cells were isolated from excisional biopsies from patients with suspected B-cell lymphomas, in accordance with a Northwestern University IRB-approved protocol (STU00208941; CSRC-1343). Samples were de-identified of all information other than eventual final diagnosis, provided to investigators approximately 1 week post excisional biopsy. A total of 7 samples were analyzed, with diagnoses of follicular lymphoma (N = 4), T-cell rich diffuse large B cell lymphoma (N = 1), diffuse large B cell lymphoma isolated from ascites fluid (N = 1), and non-GC [activated B cell (ABC)] diffuse large B cell lymphoma (N = 1). The excised tissue was washed with 1 X PBS and mechanically dissociated using two 18-G needles. The solution was then passed through a 70-micron filter, washed with 1 X PBS and centrifuged at 400 × g for 5 minutes at room temperature. The cells were re-suspended in RPMI 1640 with L-glutamine and 25mM HEPES containing 10% fetal bovine serum and 1% PenStrep. Cells were cultured at 1 × 10^6^ cells/ mL for 1 to 2 days. Following culture, CellSep Human CD19^+^ selection kit (Stem Cell Technologies) was used to enrich for CD19+ cells. Flow cytometry was used to quantify SCARB1 and CD19 expression. Data were analyzed using the FCS Express software.

To quantify cell death, CD19+ enriched primary lymphoma cells were cultured in the presence of PBS, human HDL, or HDL NPs for 72 hours. The cells were then collected, stained with Annexin V-FITC and propidium iodide (PI) (Invitrogen), and run on the BD LSR II flow cytometer (BD Biosciences). Data were analyzed with the FCS Express software. The total dead cells were analyzed as the total Annexin V-positive cell population, in both the PI positive and negative gates (Annexin V-FITC^+^/PI-) and (Annexin V-FITC^+^/PI^+^) cells. To quantify ferroptosis in primary DLBCL cells, CD19+ enriched cells were cultured in the presence of PBS, human HDL (100nM) or HDL NPs (100nM), in the presence or absence of ferrostatin-1 (final concentration of 1µM). Viability was quantified as described above. Lipid peroxide accumulation was measured using C11-BODIPY, as described above.

### Statistical Analyses

Each in vitro cell line experiment was repeated 3 times. Based on our previous experience with the SUDHL4 xenograft model, three mice were used per group. For the primary lymphoma sample assays, each sample was run once, with N = 4 or more per treatment group. One-way ANOVAs and student’s t-test were used to determine statistical significance, where appropriate. All statistical analyses were calculated using the GraphPad Prism software.

## RESULTS

### HDL NPs Bind to SCARB1 in Lymphoma

We previously demonstrated that Ramos and SUDHL4 cells, which are well studied models of Burkitt’s lymphoma (BL) and germinal center DLBCL (GC DLBCL) respectively, highly express SCARB1 (ref). Also, data showed that HDL NPs target SCARB1 in SUDHL4 and Ramos cells, which resulted in cellular cholesterol depletion and profound in vitro and in vivo cell death (21,24). Here, we verified the requirement of SCARB1 as a target of HDL NP in these lymphoma cells using an anti-SCARB1 blocking antibody and fluorescently labeled HDL NPs. HDL NPs were fluorescently labeled using the intercalating dye DiI, as previously described (21,37). Ramos and SUDHL4 cells were treated with DiI HDL NPs for 2 hours followed by flow cytometric analysis. DiI HDL NP treatment increased the fluorescent signal in both cell lines (**Supplementary Figure 1a, b**), which was reduced by co-treatment with the SCARB1 blocking antibody (1:100 dilution) but not by an isotype control antibody (**Supplementary Figure 1a, b**). DiI HDL NP treatment of Jurkat cells, a SCARB1-T cell leukemia/lymphoma cell line, did not result in increased fluorescent signal over baseline (**Supplementary Figure 1c**), confirming that the HDL NPs target SCARB1 in B cell lymphoma. We next investigated whether other BL and GC DLBCL cell lines expressed SCARB1 and if they were sensitive to HDL NP-induced cell death. The BL cell lines Raji, Daudi, Namalwa and the GC DLBCL cell line SUDHL6 all expressed SCARB1 (**Supplementary Figure 2a**). HDL NP treatment potently induced cell death in all SCARB1^+^ B cell lymphoma cell lines, while having no effect on the SCARB1-Jurkat cell line (**Supplementary Figure 2b**). Inhibition of HDL NP binding to SCARB1 using an inverse serial dilution of the SCARB1 blocking antibody protected all SCARB1^+^ cell lines from HDL NP-induced cell death in a dose-dependent manner, and the SCARB1 blocking antibody did not result in cell death when added in isolation (**Supplementary Figure 2c-h**). The isotype control antibody had no effect on HDL NP-induced cell death (data not shown).

**Figure 1:**
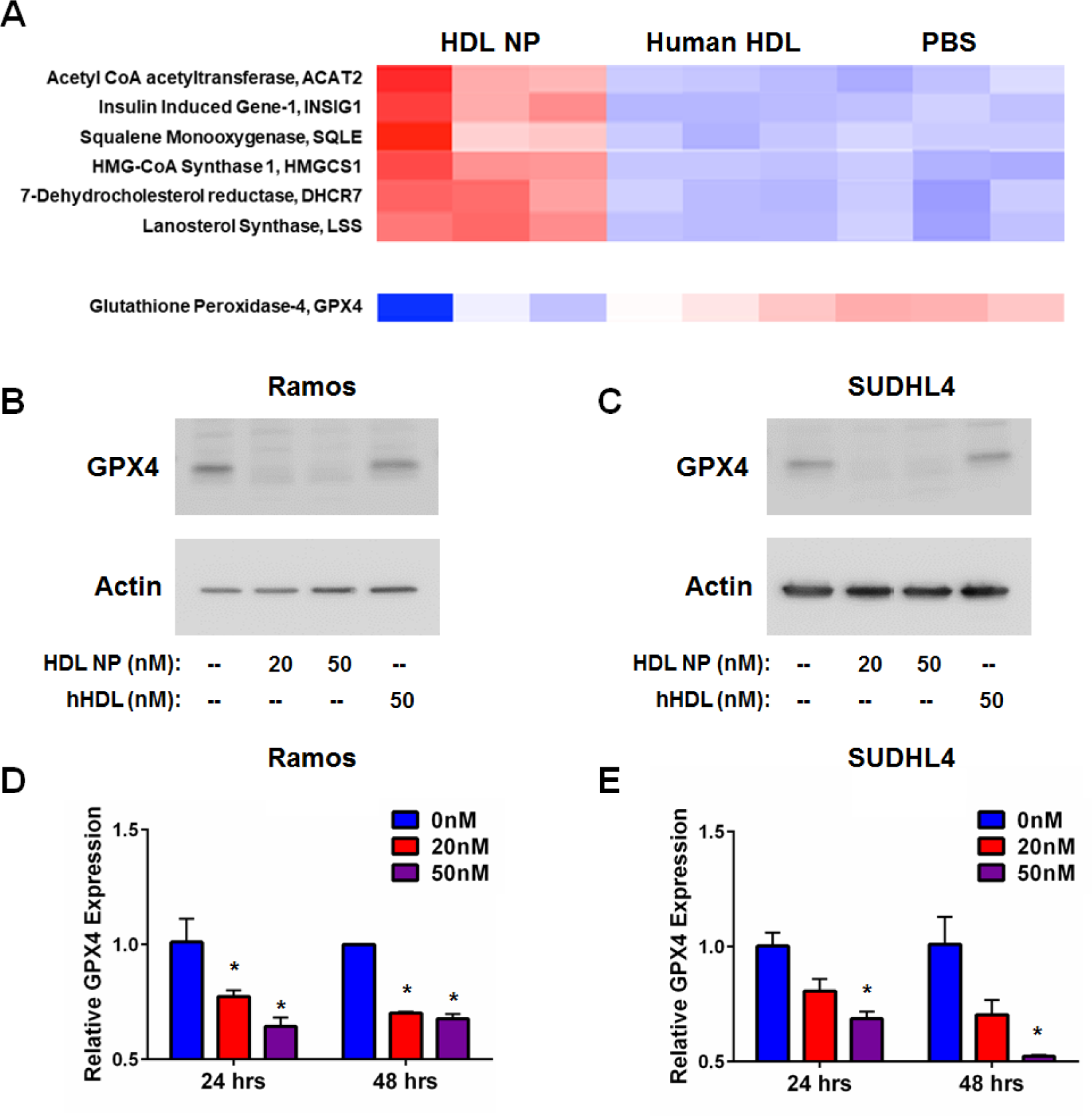
HDL NPs Increase Expression of *De Novo* Cholesterol Synthesis Genes and Reduce Expression of GPX4. A. Select gene microarray results from Ramos cells treated with HDL NPs or human HDL (hHDL) for 48 hours. B, C. Western blot analysis of GPX4 expression in Ramos (B) and SUDHL4 (C) cells treated with HDL NPs or hHDL for 48 hours. β-actin was used as a loading control. D, E. RT-qPCR analysis for GPX4 expression in Ramos (D) and SUDHL4 (E) cells treated with HDL NPs for 24 or 48 hours. *p < 0.05 vs 0nM.

**Figure 2:**
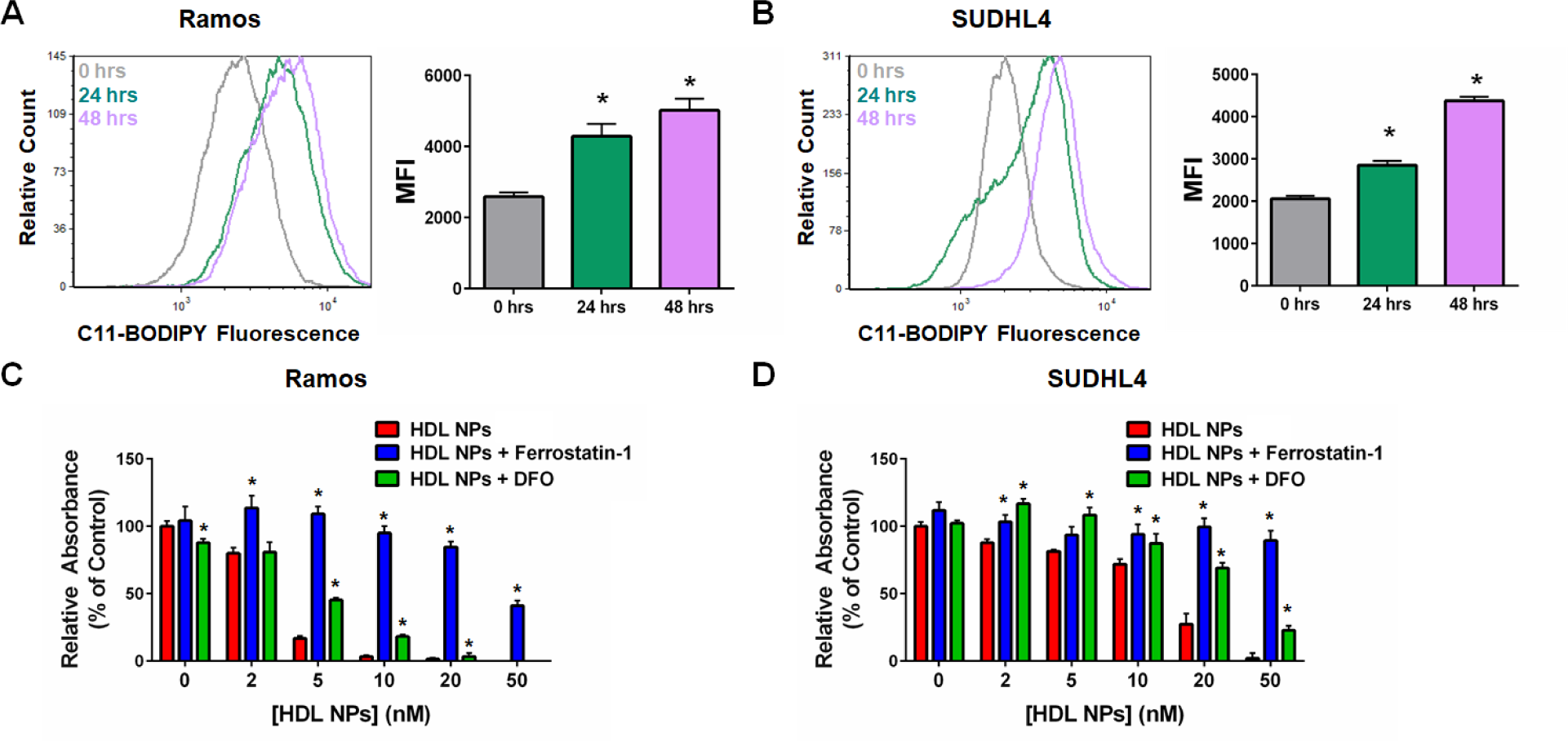
HDL NPs Induce Ferroptosis in B Cell Lymphoma Cells. A, B. Flow cytometric analysis of C11-BODIPY fluorescence in Ramos (A) and SUDHL4 (B) cells treated with HDL NPs (50nM) over time. *p < 0.05 vs. 0 hrs. C, D. Cell viability (MTS) assays for Ramos (C) and SUDHL4 (D) cells treated with HDL NPs, Ferrostatin-1 (1µM) and/ or DFO (1µM) for 72 hours. *p < 0.05 vs HDL NPs.

### HDL NP Binding to SCARB1 Increases De Novo Cholesterol Biosynthesis and Reduces GPX4 Expression in Lymphoma

We performed an unbiased gene array study in Ramos cells to broadly screen for changes in gene expression that may explain increased cell death induced by HDL NP. As a control, we treated cells with either PBS or an equimolar concentration of cholesterol-rich human HDL. We show that HDL NP treatment induces a robust increase in the expression of genes involved in de novo cholesterol biosynthesis, as anticipated (**Figure 1a**; **Supplementary Figure 3**). In addition, importantly, we found that *GPX4* was suppressed upon HDL NP treatment (**Figure 1a**; **Supplementary Figure 3**).

**Figure 3:**
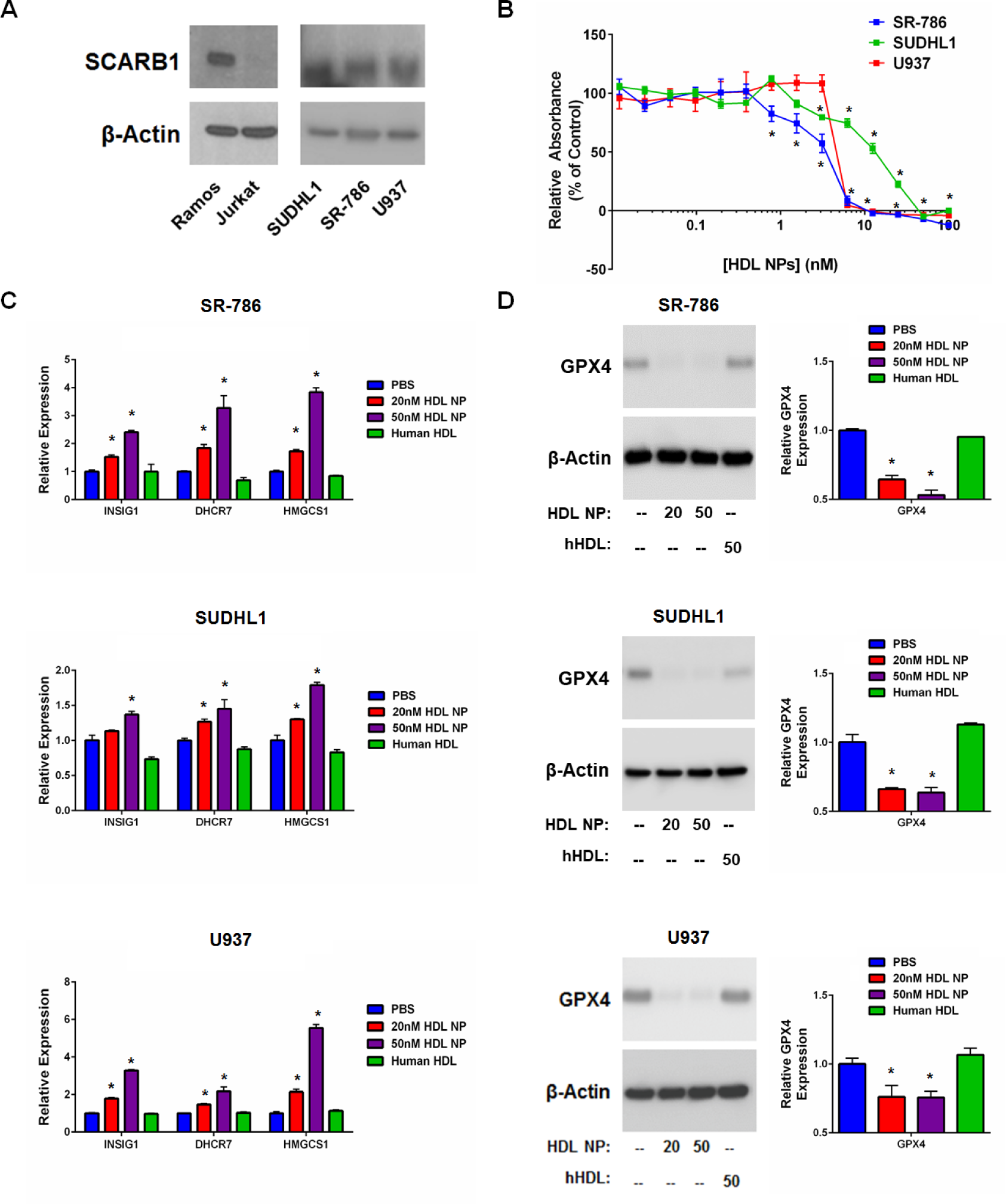
Cholesterol Auxotrophic Cell Lines Express SCARB1 and HDL NP Treatment Reduces Cell Viability and Alters Gene Expression. A. Western blot analysis of SCARB1 expression in SUDHL1, SR-786 and U937 cells. Ramos and Jurkat represent positive and negative controls for SCARB1 expression, respectively. β-actin was used as a loading control. B. Cell death (MTS) assay of SR-786, SUDHL1 and U937 cells treated with HDL NPs for 120 hours. *p < 0.05 vs PBS control. C. RT-qPCR analysis of cholesterol biosynthesis genes *INSIG1*, *DHCR7* and *HMGCS1* in SR-786, SUDHL1 and U937 cells treated with HDL NPs (20nM, 50nM) or human HDL (50nM) for 72 hours. *p < 0.05 vs PBS control. D. Western blot (left) and RT-qPCR (right) analysis of GPX4 expression in SR-786, SUDHL1 and U937 cells following 72-hour treatment with HDL NPs (20nM, 50nM) or human HDL (50nM). *p < 0.05 vs PBS control.

We confirmed decreased GPX4 expression using western blot analysis and conventional RT-qPCR. Ramos and SUDHL4 cells were treated with HDL NPs for up to 48 hours. We found that HDL NP treatment profoundly reduced expression of GPX4 in both cell lines relative to PBS control at both the protein (**Figure 1b, c**) and mRNA level (**Figure 1d, e**). By contrast, treatment with an equimolar concentration of cholesterol-rich HDL did not alter GPX4 protein or gene expression (**Figure 1a, b, c**).

### HDL NP Induces Ferroptosis in B Cell Lymphoma Cell Lines

Stockwell et al. (10), proposed two metrics to distinguish ferroptosis from apoptosis and other forms of cell death: 1) Cell death correlates with an increase in oxidized membrane lipids quantified by using C11-BODIPY, a lipophilic fluorescent dye that has a unique spectral signature when oxidized and is used to measure lipid peroxidation, and flow cytometry; and 2) Cell death can be reduced by addition of a lipophilic antioxidant (e.g. ferrostatin-1) or an iron chelator [e.g. deferoxamine (DFO)]. Using these parameters, we investigated whether HDL NPs induced ferroptosis in Ramos and SUDHL4 cells. In both cell lines, HDL NP treatment led to a dose-dependent increase in C11-BODIPY signal over time (**Figure 2a, b**). Next, Ramos and SUDHL4 cells were cultured with HDL NPs in the presence of either ferrostatin-1 or DFO and assayed for cell viability. Addition of ferrostatin-1 and DFO significantly inhibited HDL NP induced cell death in Ramos (**Figure 2c**) and SUDHL4 (**Figure 2d**) cells. Taken together, these data demonstrate that HDL NPs induce ferroptosis in Ramos and SUDHL4 cells.

### HDL NP Induces Ferroptosis in Cholesterol Auxotrophic Lymphoma Cell Lines

Recently, a number of cell lines were found to be auxotrophic for cholesterol, including the cell lines SR-786 (ALK^+^ ALCL), SUDHL1 (ALK^+^ ALCL), and U937 (isolated from histiocytic lymphoma, but of myeloid lineage), among others (16). The ALK^+^ ALCL cells were identified based upon reduced viability when cultured in lipoprotein deficient serum, and the cell death phenotype was rescued by addition of cholesterol-rich low-density lipoprotein (LDL) or free cholesterol. However, because cells can uptake cholesterol by cholesterol-rich HDL binding to SCARB1, we studied expression of SCARB1 in ALK^+^ ALCL and U937 cells. Data reveal SCARB1 expression in SR-786, SUDHL1 and U937 cells (**Figure 3a**). Furthermore, treatment of each of the cell lines with HDL NPs potently induced cell death (**Figure 3b**). Analyses of the following select de novo cholesterol biosynthesis genes in SR-786, SUDHL1 and U937 cells after HDL NP treatment demonstrated increased expression: INSIG1 (insulin induced gene-1), DHCR7 (dehydrocholesterol reductase 7) and HMGCS1 (HMG-CoA synthase 1) (**Figure 3c**). Similar to the B cell lymphoma cell lines, HDL NP treatment potently reduced GPX4 expression at both the RNA and protein levels (**Figure 3d**).

Corresponding to the decrease in GPX4 expression, HDL NPs induced accumulation of lipid peroxides in SR-786, SUDHL1 and U937 cells, as measured by C11-BODIPY flow cytometry (**Figure 4a, b, c**). Addition of the ferroptosis inhibitors ferrostatin-1 and DFO rescued SR-786, SUDHL1 and U937 cells from HDL NP induced cell death (**Figure 4d, e, f**), confirming cell death by ferroptosis. Taken together, these data indicate that inhibition of cholesterol ester uptake by HDL NP binding to SCARB1 induces ferroptosis in previously identified cholesterol auxotrophic cell lines.

**Figure 4:**
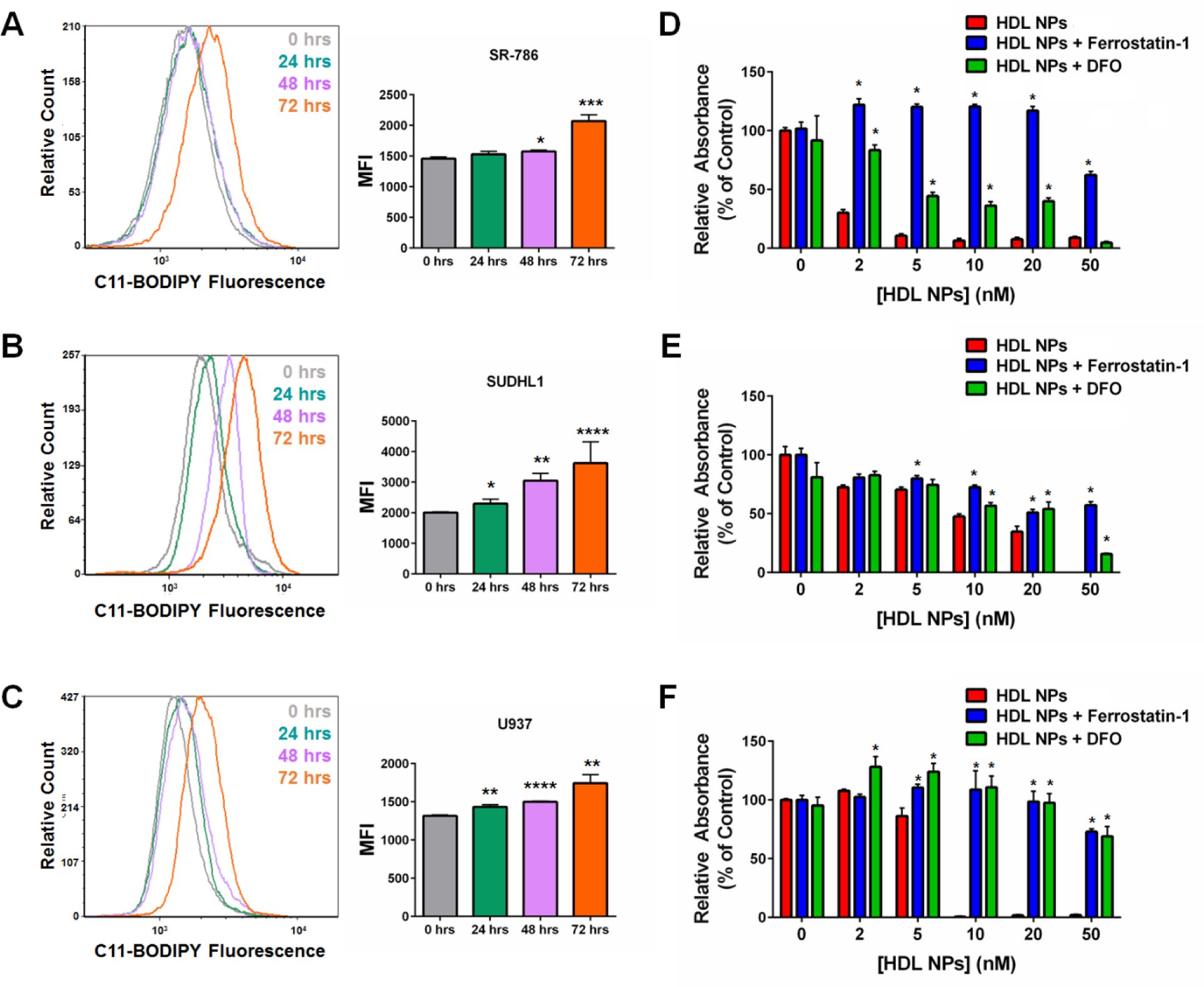
HDL NPs Induce Ferroptosis in Cholesterol Auxotrophic Cell Ines. A-C. Flow cytometric analysis for C11-BODIPY fluorescence in SR-786 (A), SUDHL1 (B) and U937 (C) cells treated with HDL NPs (50nM) over 72 hours. *p < 0.05 vs 0 hrs. **p < 0.01 vs. 0 hrs. ***p < 0.005 vs. 0 hrs. ****p < 0.001 vs. 0 hrs. D-F. Cell death (MTS) assays for SR-786 (D), SUDHL1 (E) and U937 (F) cells treated with HDL NPs, Ferrostatin-1 (1µM) and/ or DFO (1µM) for 120 hours. *p < 0.05 vs HDL NPs.

### HDL NP Induces Ferroptosis In Vivo

We previously reported that HDL NPs specifically target and significantly reduce tumor burden in xenograft models using SUDHL4 and Ramos cells (21,24). To determine if systemic HDL NP treatment reduces GPX4 expression and increases lipid peroxide accumulation in tumor cells in vivo, we established SUDHL4 tumor xenografts (~100mm^3^ in volume) in SCID-beige mice. The mice were then treated with PBS or HDL NPs (100μl of 1μM HDL NP, 3X/ week for 1 week, i.v.). Following treatment, tumors were resected and GPX4 expression and lipid peroxide accumulation were quantified by RT-qPCR and C11-BODIPY staining, respectively. HDL NP treatment led to a down-regulation of GPX4 as measured by RT-qPCR compared with PBS controls (**Figure 5a**), which correlated with an increase in membrane lipid peroxide accumulation (**Figure 5b**). No adverse side effects were observed after systemic administration of HDL NPs. These data show that HDL NPs induce molecular changes consistent with ferroptosis in the SUDHL4 flank tumor xenograft model of lymphoma previously demonstrated to be sensitive to HDL NP therapy (21).

**Figure 5.**
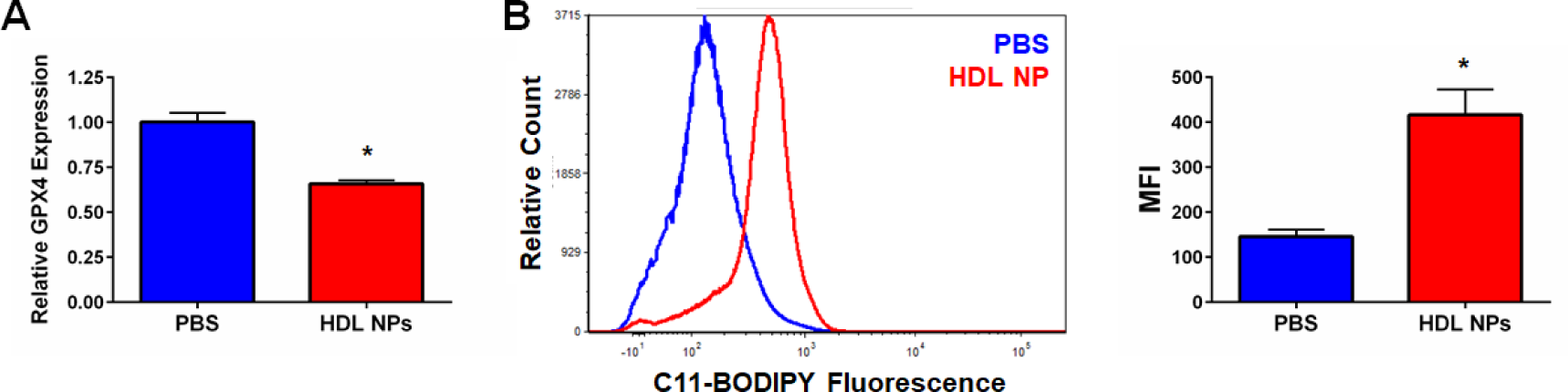
A. RT-qPCR analysis for GPX4 from SUDHL4 tumor xenografts isolated following HDL NP treatment. *p < 0.05 vs PBS control. B. Flow cytometric analysis of C11-BODIPY fluorescence in SUDHL4 tumor xenograft cells following HDL NP treatment. LEFT-Flow histogram displaying C11-BODIPY fluorescence. RIGHT-Median fluorescent intensity (MFI) graph. *p < 0.05 vs PBS control.

### In Vivo SCARB1 Expression From Archival Patient Specimens

Our in vitro data suggest that SCARB1 is a useful biomarker to identify patients with lymphoma where this targeted approach might be successful. We have previously reported, using microarray data from a cohort of DLBCL patients from a single institution, that SCARB1 expression was increased in DLBCL compared with normal naïve and memory B cells (24). In order to confirm SCARB1 expression from archival primary tissue, we performed IHC staining of formalin fixed paraffin embedded DLBCL patient samples (**Supplementary Figure 4a**). In addition, SCARB1 expression in follicular lymphoma (FL) was also investigated (**Supplementary Figure 4a**). Normal liver and thymus tissue were used as positive and negative controls, respectively. SCARB1 expression was observed in representative samples of both DLBCL and FL samples providing evidence that this is a viable target in patients (**Supplementary Figure 4a**).

### HDL NP Induces Cell Death in Primary Lymphoma Cells Obtained from Patients with Lymphoma

To best determine potential clinical relevance, we investigated expression of SCARB1 and whether HDL NPs induce cell death in primary B cell lymphoma cells derived from fresh patient tissue. Primary cells were isolated from patients with a suspected diagnosis of lymphoma after obtaining informed consent under a NU IRB-approved protocol (STU00208941; CSRC-1343). Ultimately, patients were found to have the following diagnoses: FL (**Figure 6a-d**), large cell lymphoma (T-cell rich B cell lymphoma; **Figure 6e**), DLBCL (isolated from ascites fluid; **Figure 6f**), and non-GC DLBCL (**Supplementary Figure 4b**). Cells were selected for CD19 expression to enrich the biopsy tissue samples for lymphoma cells. This resulted in a mixed population of normal B cells and B cell lymphoma cells. As we have previously shown, normal B cells do not express SCARB1 (24). Accordingly, flow cytometric analysis of the patient samples demonstrated varying levels of SCARB1 expression, as mentioned, likely the result of the mix of normal and malignant B cells (**Figure 6a-f**; **Supplementary Figure 4b**).

All of the primary patient samples were incubated with HDL NP and assayed for cell death. Cell death as a result of HDL NP exposure varied among the patient samples depending upon the lymphoma subtype and relative level of SCARB1 expression. HDL NPs dose-dependently induced cell death in the four FLs, T-cell rich B cell lymphoma, and ascites-isolated DLBCL patient samples (**Figure 6a-f**; **Supplementary Figure 4c**). Addition of human HDL had either no effect or significantly rescued the cells from cell death in culture (**Figure 6a-f**; **Supplementary Figure 4c**). In the non-GC (ABC) DLBCL sample, HDL NPs did not induce cell death at the concentrations tested (10 and 50nM; **Supplementary Figure 4b**) even with strong SCARB1 expression. This finding is consistent with our prior published work using HDL NP as monotherapy for ABC DLBCL 21. Collectively, these data confirm that HDL NP efficacy against B cell lymphoma cell lines can be replicated in primary human SCARB1^+^ B cell lymphoma cells ex vivo.

**Figure 6.**
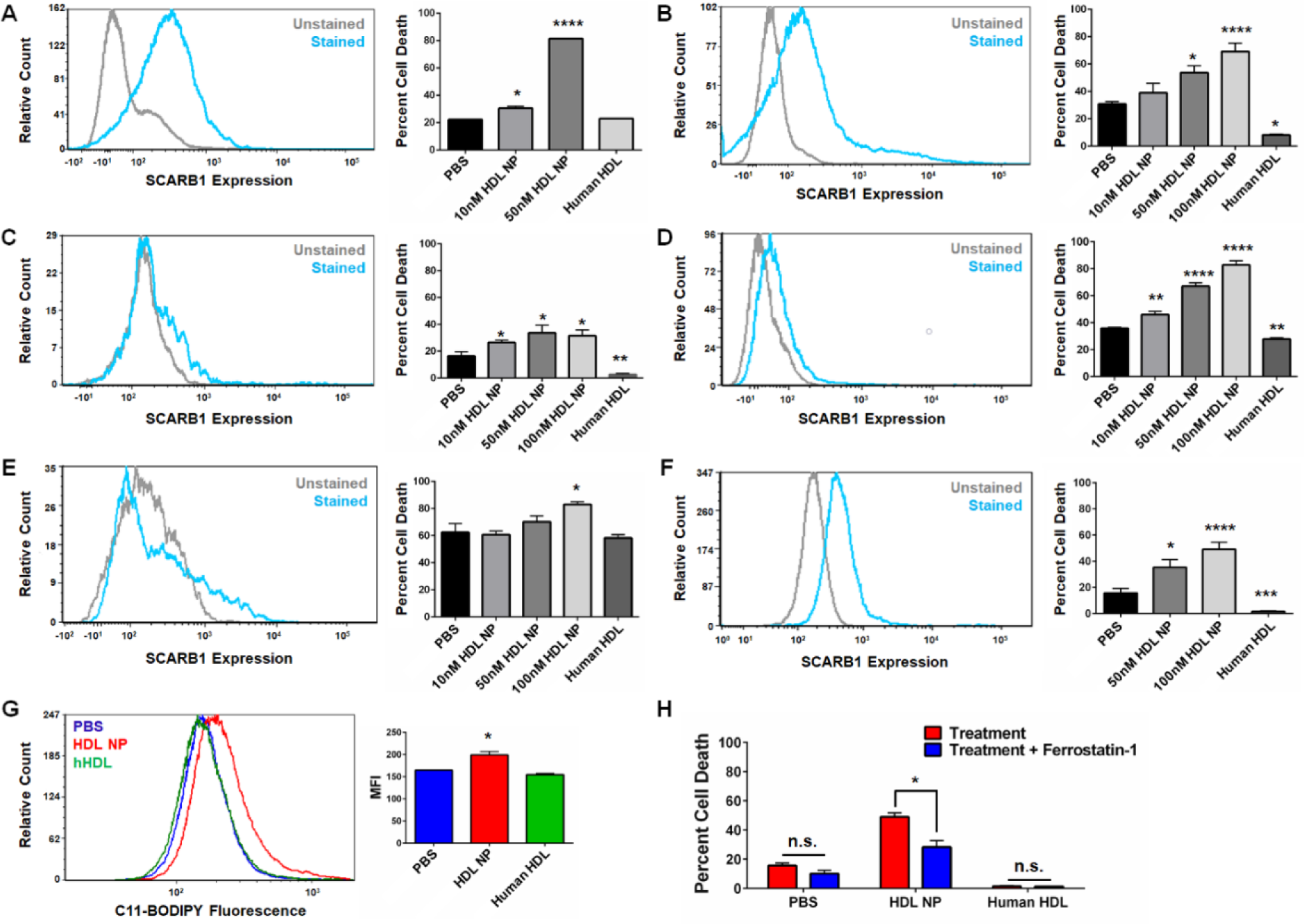
HDL NPs Induce Cell Death in Primary B Cell Lymphoma Cells Obtained From Patients. A-F. SCARB1 expression (flow cytometry, LEFT) and viability (Annexin V/ PI staining, RIGHT) of primary B cell lymphoma cells isolated from patients with follicular lymphoma and large B cell lymphoma, and treated with HDL NPs or human HDL for 72 hours (A-E) or 120 hours (F). A-D. Follicular lymphoma. E. T-cell rich large B cell lymphoma. F. DLBCL isolated from ascites fluid. *p < 0.05 vs. PBS. **p < 0.01 vs. PBS. ***p < 0.001 vs. PBS. ****p < 0.0001 vs. PBS. G. Cell death assay (Annexin V/ PI staining) for DLBCL cells from ascites fluid treated with PBS, HDL NPs (100nM) or human HDL (100nM), with or without Ferrostatin-1 (1µM) for 120 hours. *p < 0.05 vs Treatment. H. Flow cytometry for C11-BODIPY fluorescence in DLBCL from ascites fluid following treatment with PBS, HDL NPs (100nM) or human HDL (100nM) for 72 hours. *p < 0.05 vs PBS control.

To quantify if HDL NPs induce ferroptosis in the primary B cell lymphoma cells, DLBCL cells isolated from ascites fluid were assayed for lipid peroxide accumulation and viability following treatment with HDL NPs and/ or ferrostatin-1. HDL NP treatment led to an increase in measured lipid peroxide accumulation, while human HDL treatment did not (**Figure 6g**). Correspondingly, addition of ferrostatin-1 reduced HDL NP induced cell death (**Figure 6h**). Taken together, these results demonstrate that the ability of HDL NPs to induce ferroptosis in malignant cells is not restricted to immortalized cell lines. HDL NPs induce ferroptosis in primary lymphoma cells obtained from patients with lymphoma.

## DISCUSSION

The cholesterol-poor HDL NP targets SCARB1 in cholesterol uptake dependent lymphoma cells. HDL NP binding to SCARB1 results in a switch from a baseline dependence on cholesterol uptake and high GPX4 expression, to one favoring *de novo* cholesterol biosynthesis, which is accompanied by reduced expression of GPX4. As GPX4 is absolutely required by the cancer cell to reduce the burden of membrane lipid peroxides, this metabolic switch leaves the cancer cell particularly vulnerable. Accordingly, we show an increase in the accumulation of oxidized membrane lipids and cell death through a mechanism consistent with ferroptosis.

Our data demonstrate that gene and protein expression of GPX4 is downregulated after HDL NP exposure, and that this is likely mediated through the high-affinity receptor for cholesterol-rich high-density lipoproteins, SCARB1 . Gene array and RT-qPCR data support that HDL NPs mediate reduced levels of GPX4 by reducing transcription. We have shown that lymphoma cell cholesterol depletion increases the activation of SREBP-1a, which increases *de novo* cholesterol biosynthesis (21). SREBP-1a has been reported as a negative regulator of *GPX4* expression (38,39). While our data support this notion, more studies are required in order to make definitive conclusions regarding this apparent reciprocal relationship. Furthermore, our western blot data show a profound reduction in GPX4 which suggest post-translational mechanisms that would further reduce GPX4. Certainly, in this regard, there is precedent in the literature for lipid metabolism, in general, and intermediate metabolites in cholesterol biosynthesis, in particular, involved in regulating GPX4 stability (16). Interestingly, inhibiting de novo cholesterol biosynthesis using statins did not reduce GPX4 expression or induce ferroptosis (40), suggesting that manipulation of *de novo* cholesterol biosynthesis is unable to replicate the effects of HDL NP treatment. Further studies are required to better understand mechanisms through which the HDL NP therapy may regulate GPX4 stability.

Pathways involving intermediates in the cholesterol biosynthesis pathway are interesting in the context of the ALK^+^ ALCL (SR-786, SUDHL1) and U937 cell lines because of their shared inability to synthesize cholesterol due to enzymatic blockade induced by hypermethylation or mutation, respectively. Our data show that HDL NP treatment increased expression of *de novo* cholesterol synthesis genes and reduced expression of *GPX4.* In theory, this could serve to even more drastically increase intermediates in the cholesterol biosynthesis pathway that may serve an antioxidant function, but would only be effective at preventing ferroptosis in the presence GPX4. While there have been significant contributions with regard to understanding intermediates in the cholesterol biosynthesis pathway and their impact on GPX4 and ferroptosis (8,10,11,40), more work is needed in this regard. The HDL NPs provide a unique tool for further mechanistic studies and likely have translational relevance.

We have also observed that HDL NP binding to SCARB1 results in alterations in cell membrane lipid raft microdomains (34), often sites of concentrated cell membrane second messenger signaling cascades, such as the case with BCR (35). We have demonstrated that HDL NPs synergize with inhibitors of specific receptor tyrosine kinases to induce cell death in ABC DLBCL (21). Thus, it is also possible that HDL NP targeting SCARB1 modulates important downstream second messenger signaling pathways in cholesterol uptake dependent lymphomas and that this also contributes to ferroptosis.

Our data suggest that investigation of HDL NP in cholesterol auxotrophic cell lines is warranted, despite the fact that these cells can uptake cholesterol via LDLs binding the LDLR. SCARB1 expression was measured in the three auxotrophic cell lines investigated, suggesting that both the LDLR and SCARB1 play a role in supplying the cells with cholesterol. Possible explanations for our observation of potent reduction of GPX4 and ferroptosis after treatment with HDL NP include the following: a) a reduction of cholesterol uptake through LDLR by HDL NP; b) a dependence upon both LDLR and SCARB1 for sufficient cholesterol uptake; or, c) different cellular mechanisms related to cholesterol uptake through HDL via SCARB1 (cell membrane binding) versus LDL via LDLR (particle internalization). As mentioned above, and in contrast to LDL/LDLR, HDL binding to SCARB1 has been linked to intracellular signaling pathways, including the pro-survival PI3K/AKT pathway (41). A recent report suggests that a decrease in GPX4 expression correlated with decreased phosphorylation of AKT (42). As such, it is possible that engagement of HDL NPs to SCARB1 not only prevents cholesterol influx but also disrupts membrane anchored pro-survival signaling pathways that may, ultimately, impact GPX4 expression. Regardless, targeted inhibition of cholesterol uptake by synthetic nanoparticles built upon an inert core appears to be an important target in certain cholesterol auxotrophic or cholesterol uptake dependent cancers.

In this study, and in many previous studies using the HDL NP as systemic therapy in animal models, we have noted a lack of toxic side effects (21,24,25,37,43,44). We also reported that HDL NP treatment of primary human hepatocytes and macrophages, the two most abundant normal cell types that express SCARB1, did not result in toxicity and that each of the normal cell types tightly regulated cholesterol homeostasis when treated with the HDL NP or native human HDL (24). Coupled with data demonstrating the potent toxicity of HDL NPs toward lymphoma cancer cells that have been reprogrammed to depend upon cholesterol uptake and GPX4 expression to prevent ferroptosis, our working hypothesis to explain the lack of toxicity is that normal cells do not have the same oxidative burden as the cancer cells and the normal cells are able to maintain plasticity with regard to cholesterol metabolism.

In conclusion, we report that HDL NPs target SCARB1 in cholesterol uptake and GPX4 dependent lymphoma cells revealing an apparent reciprocal oncometabolic response favoring an increase in *de novo* cholesterol biosynthesis at the expense of GPX4 expression. Ultimately, this results in the death of the cancer cells by ferroptosis while sparing normal, healthy cells. As this metabolic profile is not unique to lymphomas, it is possible that HDL NPs may have translational relevance in a range of cholesterol-addicted cancers that are sensitive to cell death by ferroptosis following GPX4 depletion.

## Author Contributions

J.S.R. synthesized HDL NPs used in all experiments, assisted in testing HDL NP efficacy against primary patient derived lymphoma cells, assisted in conducting the *in vivo* SUDHL4 tumor xenograft study, and conducted SCARB1 blocking antibody studies, HDL NP binding assays, microarray analysis and RT-qPCR verification, and *in vitro* and *in vivo* ferroptosis assays. A.Y.L. assisted in acquiring the primary patient derived lymphoma cells and testing HDL NP efficacy. K.M.M. and A.E.C. assisted in conducting the *in vivo* SUDHL4 tumor xenograft study. S.Y. assisted with cell culture work. T.T. and A.B. stained and analyzed primary patient tissue samples for SCARB1 expression by IHC. J.M., R.K. and A.C. assisted in acquiring patient tissue samples for IHC analyses. A.B. and R.K. assisted in acquiring primary patient lymphoma samples for *in vitro* efficacy experiments. J.S.R., A.Y.L., R. K., C.S.T. and L.I.G. designed the experiments, analyzed the data, and wrote the manuscript.

## Acknowledgements

This work was supported by the Robert H. Lurie Comprehensive Cancer Center Cancer Center Support Grant (RHLCCC CCSG) (LIG, CST) the National Institutes of Health/National Heart, Lung, and Blood Institute for a Vascular Surgery Scientist Training Program grant (T32HL094293) (JSR), the American Society of Hematology Research Training Award for Fellows (AYL), the Department of Defense/Air Force Office of Scientific Research (FA95501310192), the CRN Regenerative Nanomedicine Catalyst Award Program at Northwestern University, and the National Institutes of Health/National Cancer Institute (R01CA167041) (CST) and the Brookstone, Shannahan and Mander Foundations (LIG). We thank the RHLCCC Northwestern University (NU) Flow Cytometry Core Facility, the RHLCCC NU Immunobiology Center Flow Cytometry Core Facility and the RHLCCC Pathology Core.

