## Supplemental Figures 1-4 for "Targeting Scavenger Receptor Type B1 In Cholesterol-Addicted Lymphomas Abolishes Glutathione Peroxidase 4 and Results in Ferroptosis"

**Supplementary Figure 1**

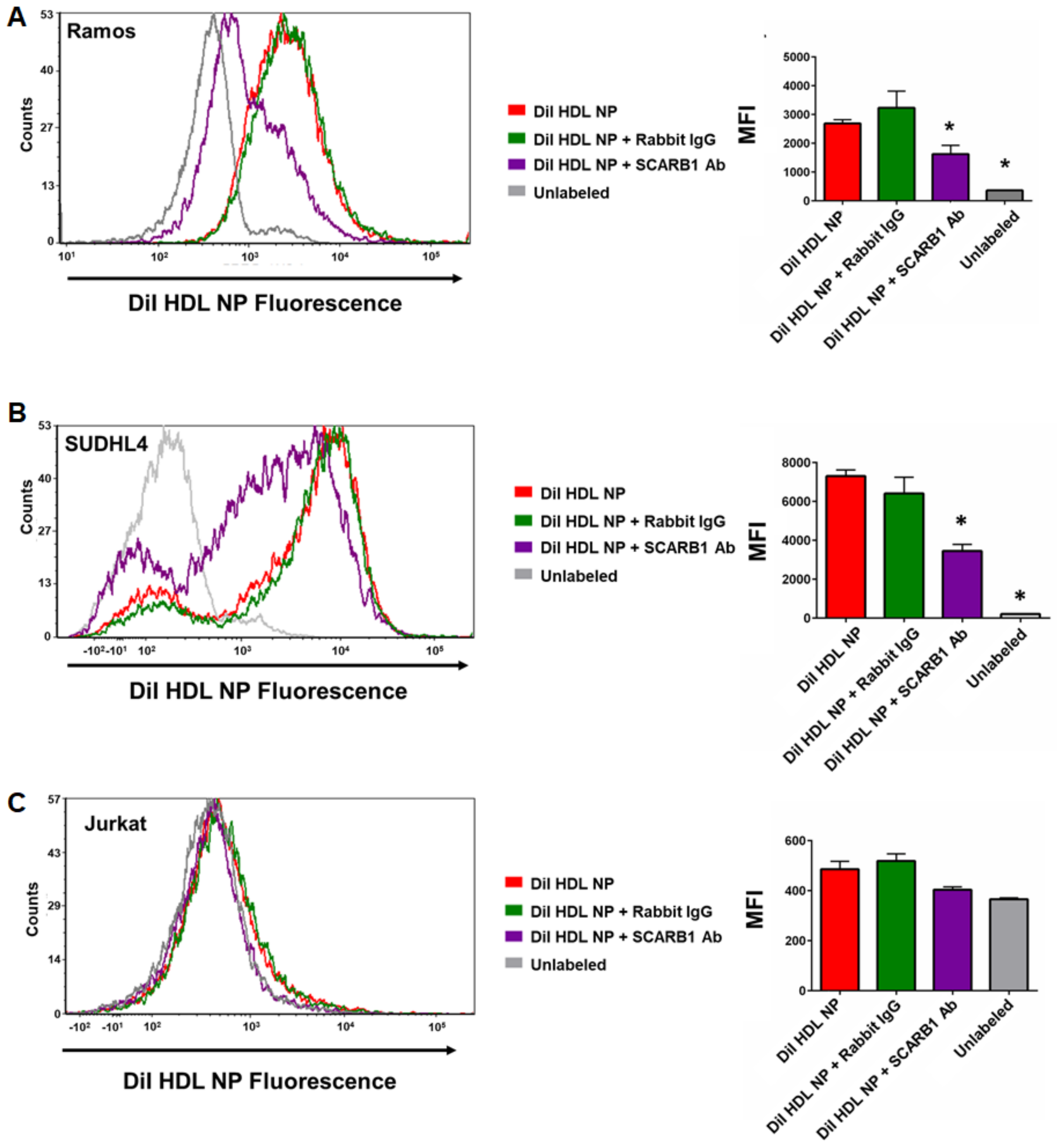


**Supplementary Figure 1: HDL NPs Bind to SCARB1 on B Cell Lymphoma Cells.** Flow cytometric analysis of the binding of DiI HDL NPs to Ramos (A), SUDHL4 (B) and Jurkat cells (C) treated with the SCARB1 blocking antibody (1:100), Rabbit IgG isotype control antibody (1:100), or untreated. Left- Representative histogram of DiI HDL NP treated cells. Right- Median fluorescent intensity. *$p<0.05$ vs. DiI HDL NPs.

**Supplementary Figure 2**

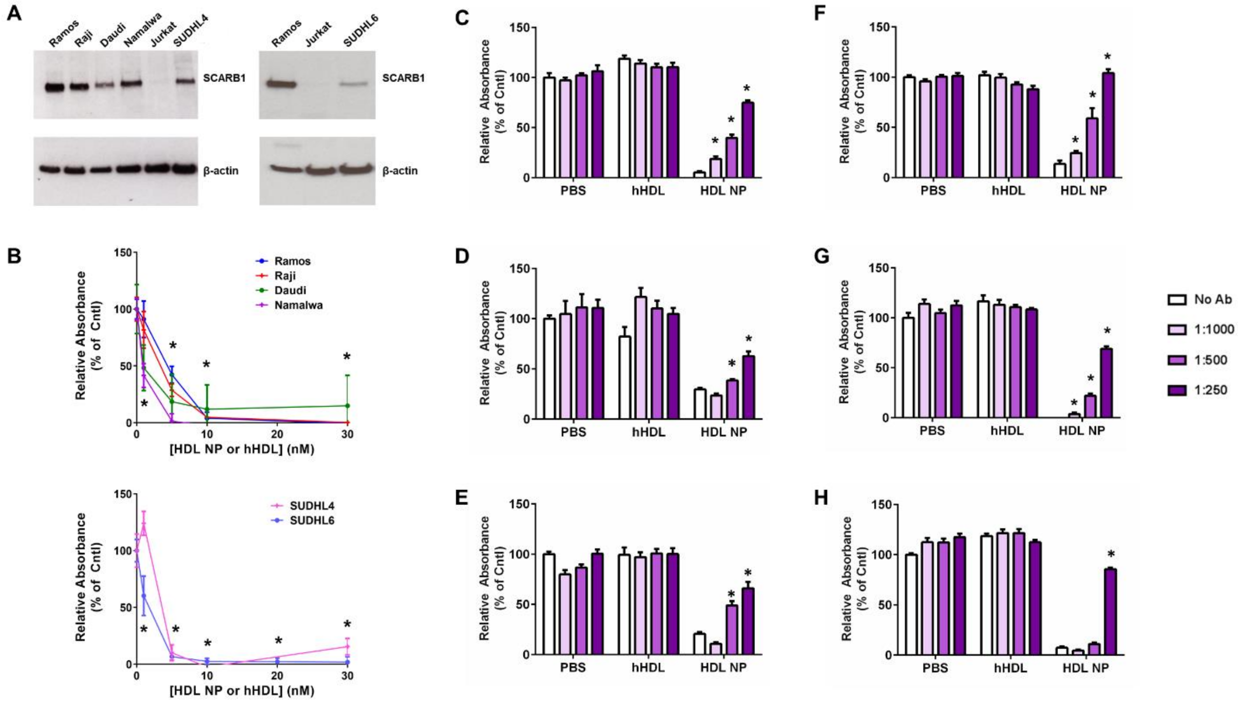


**Supplementary Figure 2: HDL NPs Induce Cell Death in GC DLBCL and BL Cell Lines Through Binding SCARB1.** A. Western blot analysis for SCARB1 in BL (Ramos, Raji, Daudi, Namalwa) and GC DLBCL (SUDHL4, SUDHL6) cell lines. The SCARB1-negative T cell leukemia/ lymphoma cell line Jurkat was used as a negative control. β-actin was used as a loading control. B. MTS assays for BL (Top) and GC DLBCL (Bottom) cell lines treated with increasing concentrations of HDL NPs. N=6 for all conditions. *$p<0.05$ vs. PBS control. C-H. Cell survival as assessed by MTS assays of SCARB1 blocking antibody treated Ramos (C), Raji (D), Daudi (E), Namalwa (F), SUDHL4 (G), and SUDHL6 (H) cells exposed to human HDL (hHDL) or HDL NPs (10nM). N=6 for all conditions. *$p<0.05$ v. No Ab.

**Supplementary Figure 3**

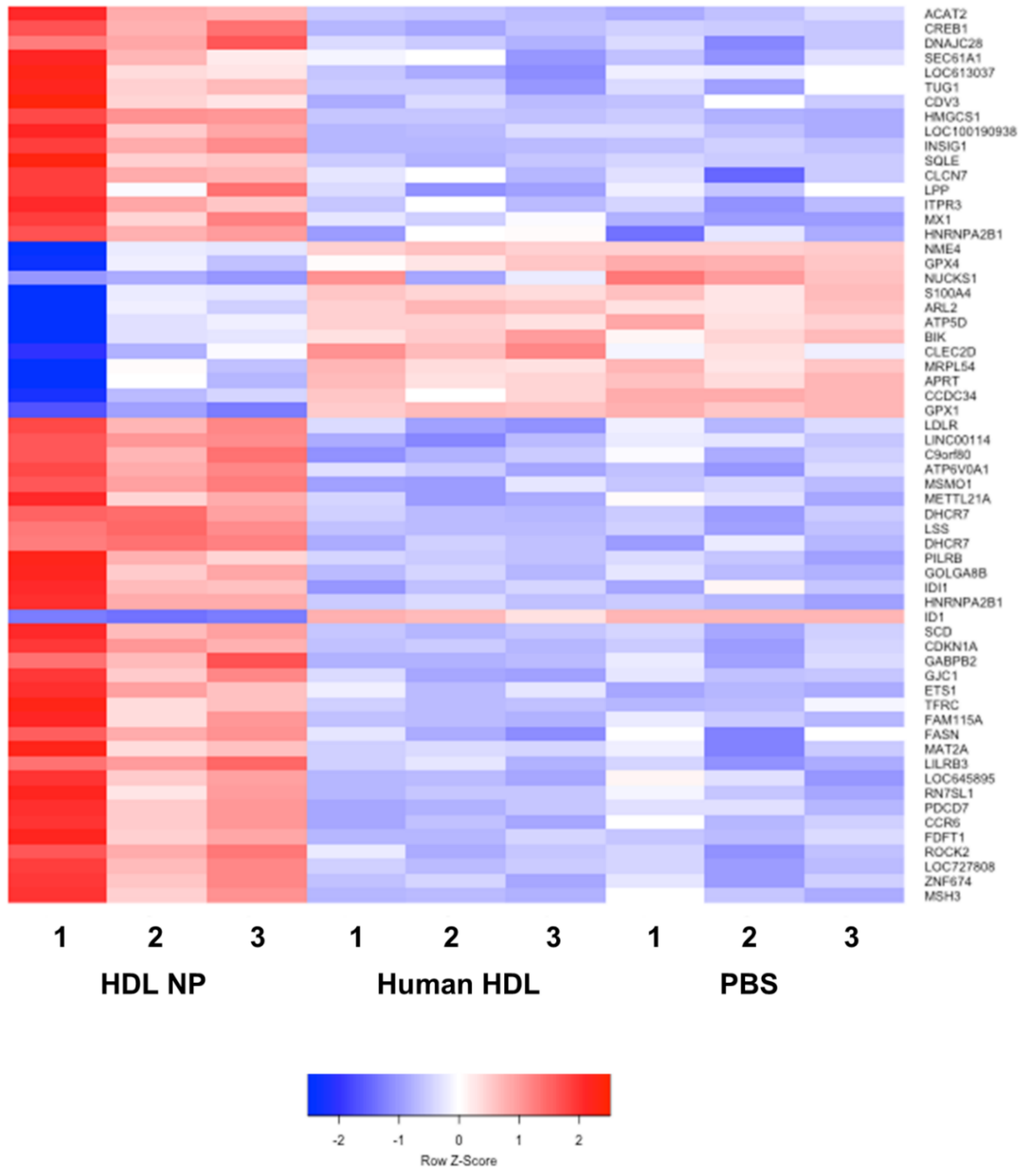


**Supplementary Figure 3: HDL NPs Alter Gene Expression in Ramos Cells.** Ramos cells were treated with PBS, human HDL (40nM) or HDL NPs (40nM) for 48 hours prior to RNA extraction for analysis by Illumina's HT-12 microarray. All genes listed had a fold change of >1.5 or <-1.5 vs. PBS control, with a corresponding p value less than 0.05.

**Supplementary Figure 4**

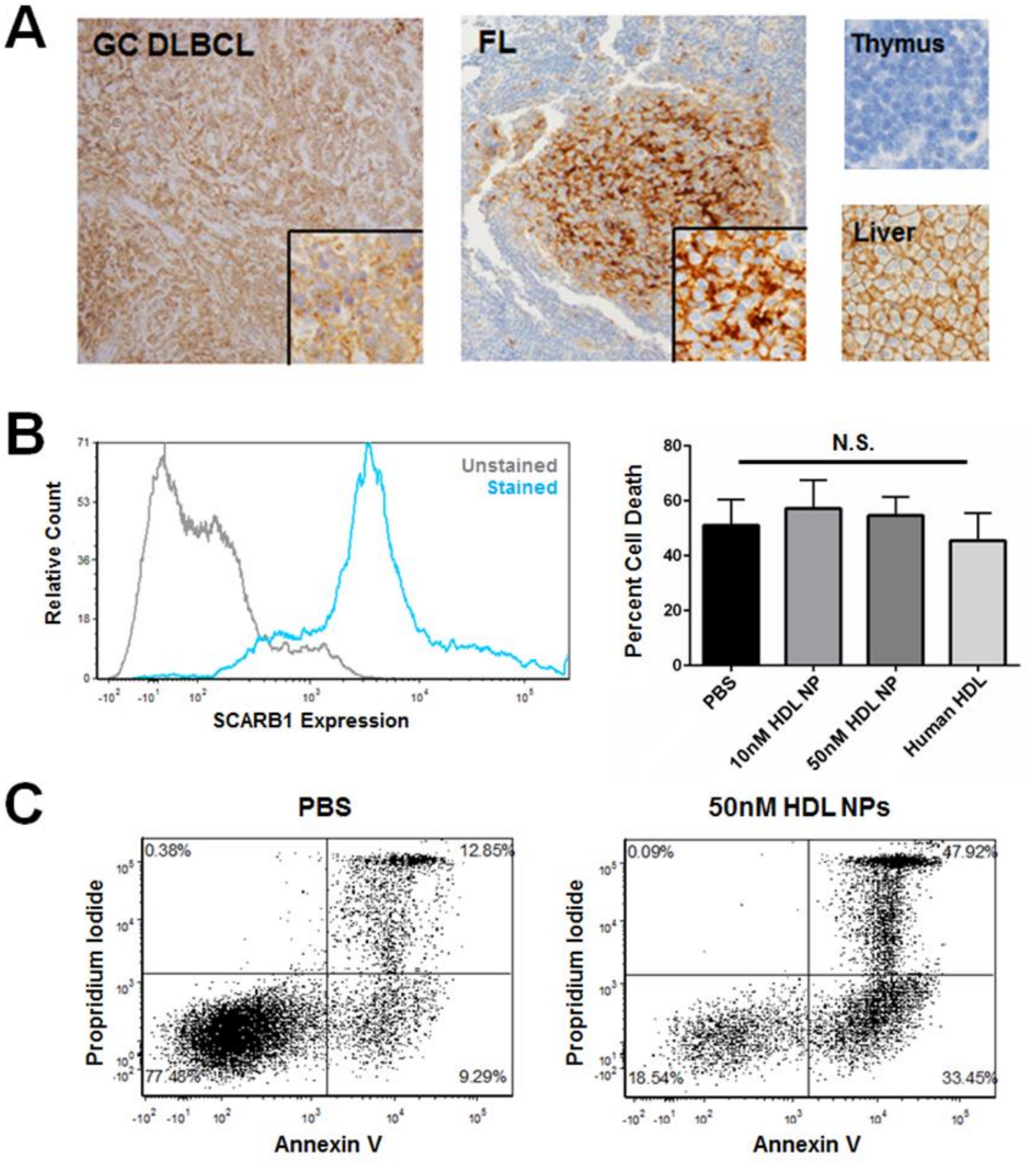


**Supplementary Figure 4: SCARB1 Expression and HDL NP Efficacy Against Primary Lymphoma Cells.** A. Formalin-fixed, paraffin embedded tissue samples, obtained at the time of diagnosis were sectioned and stained for SCARB1 (brown staining). Representative positive staining of DLBCL (N = 49; 26.5% of which displayed ≥10% of malignant cells positive for SCARB1) and FL (N = 104; 10.6% of which displayed ≥10% of malignant cells positive for SCARB1). Thymus and Liver sections (bottom) are presented as negative and positive staining controls, respectively. Images were taken at 10X magnification. Insets and control images were taken at 40X magnification. B. Flow cytometric analyses of SCARB1 and viability of HDL NP treated primary DLBCL cells isolated from a patient with non-GC (ABC) DLBCL. C. Representative dot plots of Annexin V/ PI stained primary FL cells treated with PBS or 50nM HDL NPs.
